# Temporal expectation hastens decision onset but does not affect evidence quality

**DOI:** 10.1101/2020.01.30.926337

**Authors:** Ruud L. van den Brink, Peter R. Murphy, Kobe Desender, Nicole de Ru, Sander Nieuwenhuis

## Abstract

The ability to predict the timing of forthcoming events, known as temporal expectation, has a strong impact on human information processing. Although there is growing consensus that temporal expectations enhance the speed and accuracy of perceptual decisions, it remains unclear whether they affect the decision process itself, or non-decisional (sensory / motor) processes. Here, healthy human participants (N = 21; 18 female) used predictive auditory cues to anticipate the timing of low-contrast visual stimuli they were required to detect. Modelling of the behavioral data using a prominent sequential sampling model indicated that temporal expectations speeded up non-decisional processes but had no effect on decision formation.

Electrophysiological recordings confirmed and extended this result: temporal expectations hastened the onset of a neural signature of decision formation, but had no effect on its build-up rate. Anticipatory alpha-band power was modulated by temporal expectation, and co-varied with intrinsic trial-by-trial variability in behavioral and neural signatures of the onset latency of the decision process. These findings highlight how temporal predictions optimize our interaction with unfolding sensory events.

**SIGNIFICANCE STATEMENT:** Temporal expectation enhances performance, but the locus of this effect remains debated. Here, we contrasted the two dominant accounts: enhancement through (1) expedited decision onset, or (2) an increase in the quality of sensory evidence. We manipulated expectations about the onset of a dim visual target using a temporal cueing paradigm, and probed the locus of the expectation effect with two complementary approaches: drift diffusion modeling of behavior, and estimation of the onset and progression of the decision process from a supramodal accumulation-to-bound signal in simultaneously measured EEG signals. Behavioral modeling and neural data provided strong, converging evidence for an account in which temporal expectations enhance perception by speeding up decision onset, without affecting evidence quality.

## INTRODUCTION

To efficiently process the large amount of sensory information that we constantly receive, the brain actively predicts upcoming sensory input rather than passively registering it. One way the brain achieves this is by exploiting temporal contingencies in the continuous stream of sensory input. These contingencies can be used to prepare for relevant events and optimize processing of those events. The temporal expectations shaped by these contingencies have a profound impact on perception and action, enhancing the speed and, in some cases, the accuracy of responding in a wide range of information-processing tasks (Denison, Heeger, & Carrasco, 2017; Hackley & Valle-Inclán, 2003; Niemi & Näätänen, 1981; Nobre, Correa, & Coull, 2007; Nobre & van Ede, 2018). Recent research has established that the brain expresses such temporal expectations by synchronizing oscillatory neural dynamics (power and phase) with the temporal structure of the environment (Busch, Dubois, & VanRullen, 2009; Cravo et al., 2013; Henry, Herrmann, & Obleser, 2014; Schroeder & Lakatos, 2009; Stefanics et al., 2010; van den Brink, Wynn, & Nieuwenhuis, 2014). For example, fluctuations in the amplitude of ongoing neural oscillations in the α band (9-12 Hz) closely track the time course of temporal expectations (Heideman et al., 2018; Rohenkohl & Nobre, 2011; Zanto et al., 2011). In contrast, the mechanisms through which temporal expectations enhance task performance remain unclear. The goal of the current study was to identify how temporal expectations shape perception.

Although there is a growing consensus that temporal expectations enhance the speed and/or accuracy of perceptual decisions (Correa, Lupiáñez, Madrid, & Tudela, 2006; Correa, Lupiáñez, & Tudela, 2005; Jepma, Wagenmakers, & Nieuwenhuis, 2012; Rohenkohl et al., 2012a; Rolke & Hofmann, 2007; Vangkilde, Coull, & Bundesen, 2012), different studies have arrived at different conclusions as to whether this is achieved through (1) expedited decision onset (Bausenhart et al., 2010; Jepma et al., 2012; Seibold et al., 2011), or (2) an increase in the quality of the sensory evidence (Cravo et al. 2013; Rohenkohl et al., 2012a; Vangkilde et al., 2012)—two accounts that are not mutually exclusive. Under most computational frameworks for decision-making, decision onset is determined by the duration of non-decisional processes (i.e. sensory encoding), whereas evidence quality is equivalent to the mean rate at which evidence is accumulated (i.e., decision formation; Rohenkohl et al., 2012a; Vangkilde et al., 2012).

One way to address this discrepancy around the locus of temporal expectation effects is by decomposing performance on cognitive tasks into latent information-processing parameters using sequential-sampling models (Forstmann, Ratcliff, & Wagenmakers, 2016), and examining the effects of temporal expectations on those parameters. Previous attempts to discriminate between the decision onset account and evidence quality account employed such model-based analyses of behavioral data. Rather than solely considering average RTs and error rates, model-based analyses generally consider the full distributions of RTs and their relationship with error rates, which are expected to be influenced differentially by various latent parameters, depending on task constraints such as stimulus masking or response deadlines. However, recent studies on perceptual decision-making have revealed that conclusions based on prominent sequential-sampling models can in some cases be contradicted by complementary analyses of neural signatures of decision formation (McGovern, Hayes, Kelly, & O’Connell, 2018; Spieser, Kohl, Forster, Bestmann, & Yarrow, 2018), suggesting that it is critical to corroborate insights from modelling with neural evidence. Specifically, non-invasive human EEG recordings have identified a domain-general build-to-threshold signal, the centroparietal positivity (CPP), that has been suggested to reflect decision formation via the gradual accumulation of sensory evidence (O’Connell, Shadlen, Wong-Lin, & Kelly, 2018). Importantly, measures of CPP onset latency and slope can be used to dissociate decision onset from other influences on the decision process (evidence strength and the rate of accumulation; Loughnane et al., 2016).

To address the outstanding discrepancy in the literature, we combined these modelling and electrophysiological approaches to examine the effects of temporal expectations on perceptual decision-making in a temporal cuing task, a powerful paradigm for manipulating temporal expectations (Coull & Nobre, 1998). The two approaches provided strong, converging evidence for an account in which temporal expectations enhance task performance specifically by shortening the time prior to the onset of the decision process, and not by affecting the decision process itself. In addition, time–frequency analysis of the EEG data identified peri-stimulus α-band power as a significant factor underlying the effect of temporal expectation on decision onset.

## MATERIALS AND METHODS

### Participants

A total of 30 participants took part in the study. After EEG artifact rejection and predetermined exclusion criteria (see below), a total of 21 participants remained (mean age 22.6 years old; SD 2.4; range 18-28; 18 female). All participants had normal or corrected-to-normal vision, and were free from any neurological or psychiatric disorders. Participants gave written informed consent and were compensated with €7.50 or course credit. The experiment was approved by the Leiden University Institute of Psychology Ethics Committee.

### Task

The task consisted of the detection of the appearance of a small opening (decrease in luminance, 0.19° by 0.23° visual angle) on either side of a white square box (side length = 2.34° visual angle) presented on a black background (Figure 1a). Participants reported the onset of the target stimulus (i.e., the small opening) by pressing the space bar of a computer keyboard with the index finger of their right hand, regardless of whether the target appeared on the left or the right. The target remained on the screen for a limited viewing time of one second, during which the participant was required to respond. If the participant responded after this time, the trial was counted as an error. Upon target offset, the surrounding box also disappeared briefly (50 ms) to signal the onset of a new trial. If the participant responded before target onset, the text “You responded too soon!” appeared in red and the trial was aborted. Task difficulty was manipulated by adjusting the luminance of the target stimulus. The decrease in luminance relative to the surrounding box on easy and difficult targets (28% and 20%) was determined in a pilot experiment (*N* = 6), and set such that its effect on RT was of approximately the same size as the effect of cue validity on RT for the short cue-target interval (CTI; see below). This allowed a fair dissociation between underlying model parameters and electrophysiological markers. We opted for a detection task rather than a discrimination task because temporal cuing effects on RT are consistently found for detection tasks but not discrimination tasks (Correa et al., 2004), and we intended to leverage such effects in modeling analyses and in probing the electrophysiological data for signatures of the underlying processes at play.

**Figure 1.**
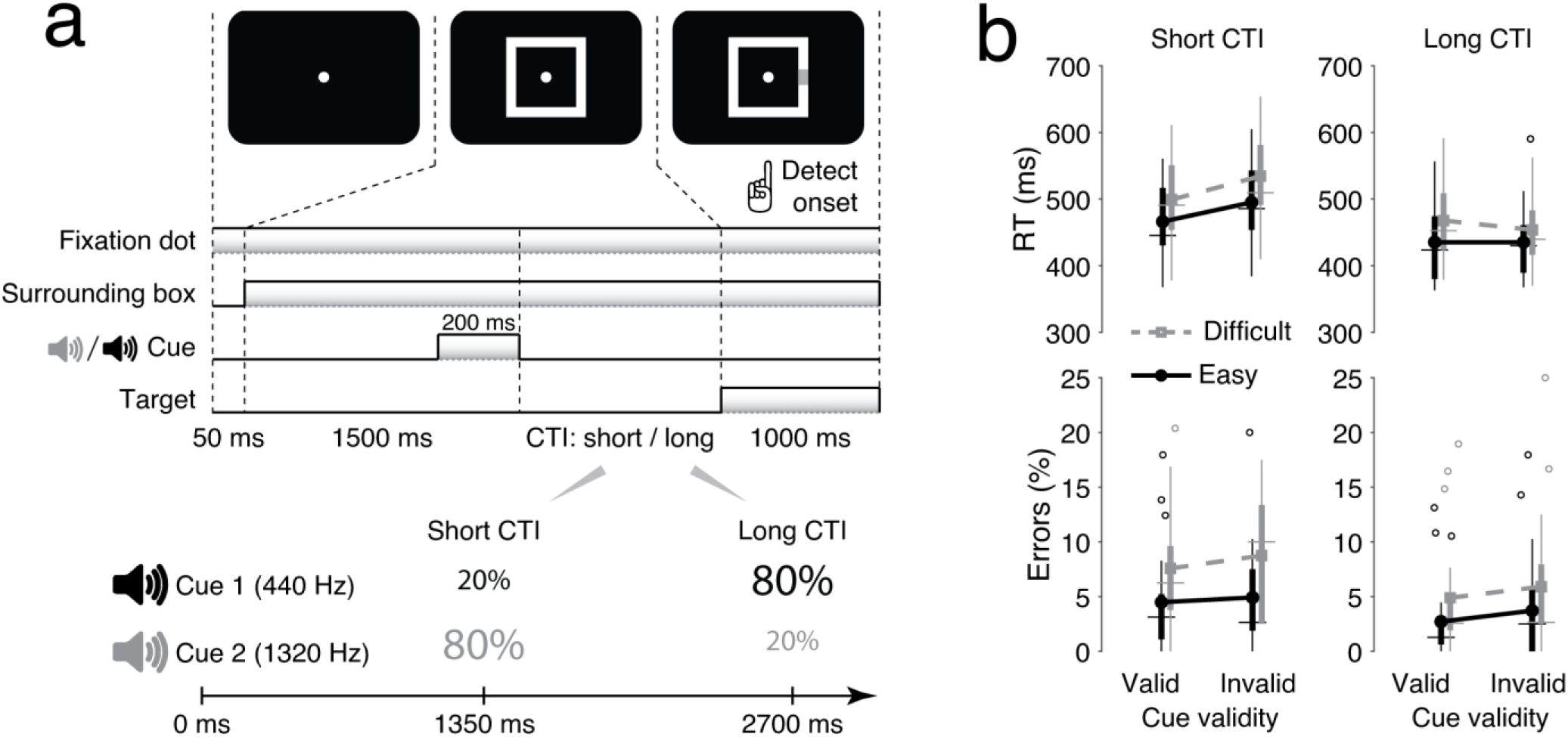
**a)** Task design, and **b)** behavioral results. The error rate includes both types of errors (false alarms and misses). CTI: cue-target interval RT: response time. Horizontal markers on the box plots indicate the median, the bottom and top edges indicate the 25th and 75th percentile, and the whiskers extend to the most extreme non-outlier data points. Outliers are shown as circles.

In order to manipulate temporal expectation about target onset, we used the temporal cueing paradigm (Coull & Nobre, 1998). Following a 1.5 s fixation interval, target onset was probabilistically cued by a brief (150 ms) auditory signal. Cue 1 (440 Hz) signaled that the target would likely appear after 2700 ms (long CTI), and cue 2 (1320 Hz) signaled that the target would likely appear after 1350 ms (short CTI). The CTIs were chosen based on prior work (Stefanics et al., 2010). Both cues had a validity of 80%, such that on 20% of the trials the target would appear after the uncued interval (invalid trials). The cues and the target were presented in different sensory modalities so that participants could optimally distinguish them. The cue was presented in the auditory domain to ensure that participants perceived the cue even when briefly losing fixation.

As is common in the temporal cueing literature, we included a small proportion (13%) of catch trials, on which no target appeared after the cue. This ensures that participants generally await target presentation at the long CTI and make few anticipatory responses (Correa et al., 2006). These trials, along with a small amount of trials (mean = 4.0 s, SD = 5.1) on which participants responded before target onset (i.e. false alarms), were excluded from all analyses, with the exception that false alarms were included to assess effects on error rate. Catch trials were excluded because they do not have an associated RT to fit (see below) and because they are unmatched to non-catch trials in terms of their hazard function for errors (i.e. compared to other trials, catch trials contain a longer time window during which a response could be made that would count as a false alarm).

Participants were instructed to maintain fixation throughout the task, to use the cue to speed up target detection, and to “respond as quickly and accurately as possible”. In total, participants performed 8 blocks of 115 trials per block (920 in total). Participants briefly practiced the task beforehand (2 blocks of 24 trials). The total duration of the task was approximately 1 hour.

### Behavioral data analysis

Effects of cue validity, CTI, and difficulty on RT were tested with a repeated-measures ANOVA in JASP version 0.9.2 (JASP Team, 2018), with cue validity (valid or invalid), CTI (short or long), and difficulty (easy or difficult) as within-participant factors. Planned paired-samples *t* tests were conducted to examine the effect of cue validity on RT for the short CTI, the effect of difficulty on RT, and differences in the size of these two effects. Because we expected the effect of cue validity for the short CTI to be approximately the same size as the effect of difficulty, we calculated a Bayes factor (using default priors) for this statistical comparison in order to estimate the evidence for the null hypothesis of no difference. Bayes factors between zero and one indicate evidence for the null hypothesis, with 1/10 ≤ BF ≤ 1/3 indicating “substantial” evidence for the null hypothesis (Wetzels & Wagenmakers, 2012).

Accuracy was analyzed using generalized mixed regression modeling, separately for the two types of possible errors on the task (misses and false alarms). This method allowed analyzing accuracy data at the single-trial level, and was necessary given that the bounded nature of accuracy scores results in inherent violations of the assumptions made by ANOVA. We fitted random intercepts for each participant; error variance caused by between-subject differences was accounted for by adding random slopes to the model. The latter was done only when this significantly increased the model fit. We used logistic linear mixed models, for which χ^2^ statistics are reported. Model fitting was done in R (R Development Core Team, 2008) using the lme4 package (Bates et al., 2015).

### Hierarchical drift diffusion modeling

We decomposed behavioral data from the target-detection task into latent parameters of the decision process using the drift diffusion model (DDM), a popular instance of sequential-sampling models of RT tasks (Forstmann, Ratcliff, & Wagenmakers, 2016; Ratcliff & McKoon, 2008). The DDM assumes that for two-alternative forced choice decisions, noisy sensory evidence is accumulated from a starting point *z*, at drift rate *v*, toward one of two decision bounds (thresholds), labeled 0 and *a*. When the accumulated evidence reaches one of the two bounds, the corresponding decision is initiated. The distance between the bounds, referred to as boundary separation, is equal to *a*. The model ascribes all non-decisional processes, including sensory encoding and response execution, to a non-decision time parameter *T*_*er*_.

Most decision-making tasks require selecting between two overt responses. However, our task is similar to a go/no-go task, in that the participants have to arbitrate between a simple go decision when the target is presented and a no-go decision when the target is not presented. As a consequence, RTs for a no-go decision cannot be empirically measured. Therefore, in line with previous studies (e.g., Gomez, Ratcliff, & Perea, 2007; Zhang et al., 2016), we assumed an implicit absorbing lower decision bound for no-go decisions and an explicit absorbing upper boundary for go decisions. We then fitted the DDM to participants’ decisions (i.e., the proportion of go and no-go decisions) as well as the distributions of RTs (for regular trials with correctly timed responses).

We fitted this DDM to behavioral data (choices and RTs), using the hierarchical Bayesian model fitting procedure implemented in the HDDM toolbox (version 0.6.1; Wiecki, Sofer, & Frank, 2013). The HDDM uses Markov-chain Monte Carlo sampling, which generates full posterior distributions over parameter estimates, quantifying not only the most likely parameter value but also uncertainty associated with each estimate. Due to the hierarchical nature of the HDDM, estimates for individual participants are constrained by group-level prior distributions. In practice, this results in more stable estimates for individual participants, especially when working with low trial numbers (Vandekerckhove, Tuerlinckx, & Lee, 2011; Wiecki et al., 2013).

For each model fit, we drew 100,000 samples from the posterior distribution. The first 10% of these were discarded as burn-in and every second sample was discarded for thinning, reducing autocorrelation in the chains. Group-level chains were visually inspected to ensure convergence, ruling out sudden jumps in the posterior and ruling out autocorrelation. Additionally, the model that is reported in the main text was run three times in order to compute Gelman-Rubin 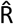 statistics (comparing within-chain and between-chain variance). We checked and confirmed that all group-level parameters had an 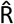 value between 0.98-1.02, suggesting convergence between these three instantiations of the same model. Because individual parameter estimates are constrained by group-level priors, data are not independent and therefore frequentist statistics cannot be used. The probability that a condition differs from another can be computed by calculating the overlap in posterior distributions. The fits of models of different complexity were compared to each other by calculating the deviance information criterion (DIC; Spiegelhalter, Best, Carlin, & van der Linde, 2002). Lower DIC values indicate that a model explains the data better, while taking model complexity into account. DIC differences larger than 10 are generally taken as strong evidence for a difference in model goodness-of-fit.

Cue validity was expected to reliably influence behavior for early, but not late targets (Correa et al., 2006). The lack of an effect for the long CTI is caused by the fact that on invalidly cued long-CTI trials, the participant can reorient temporal attention from the expected short CTI to the actual long CTI, in time for the target to appear. For this reason, we only fitted the behavioral data from trials with a short CTI. Trials on which participants failed to respond (i.e., misses, mean = 24.4 trials, SD 18.1) were considered trials on which the no-go bound was crossed. For these types of trials, RTs were set to NaN (i.e., not a number), so that the (missing) RT did not contribute to the parameter estimation (as implemented in HDDM version 0.6.1), whereas the (erroneous) decision itself did contribute, thus facilitating accurate and reliable estimates of model parameters. The trials included for fitting the DDM were thus hits (trials with an RT between target onset and the response deadline), and misses (with RT set to NaN), both in the short CTI condition only.

In the main text, we focus on a model in which both drift rate and non-decision time were free to vary as a function of cue validity, and drift rate was also free to vary as a function of difficulty. Apart from this model, we fitted and compared a variety of different models in which drift rate was always allowed to vary as a function of difficulty, and in which all possible combinations of drift rate, non-decision time and/or boundary separation were allowed to vary as a function of cue validity. Previous fits of the DDM to go/no-go tasks have allowed for the possibility of a biased starting point *z* (Gomez et al., 2007). This was possible because the tasks to which these fits were made produced a moderate number of errors (and associated RTs) on no-go trials, which were essential for appropriately constraining estimates of the *a* and *z* parameters when both were allowed to vary. In our case, there were very few such trials (i.e. behavioral responses within the temporal interval that targets could appear, but on trials when the target was not presented) and we therefore lacked the appropriate constraint to estimate *a* and *z* in the same model. For this reason, all models assumed an unbiased starting point (*z* = *a*/2; see also Ratcliff & van Dongen, 2011). We note that if the key effect of temporal expectation on behavior was in fact on starting point, in our fits this would load onto the boundary separation parameter (intuitively, if crossing of only one of the two available boundaries occurs regularly and yields measurable behavior, then a shift in starting point toward that boundary will generate identical behavior as a decrease in boundary separation of the same magnitude). However, the most complex model (in which all parameters depended on validity; Table 2) yielded no significant effect of cue validity on decision bound, making it unlikely that shifts in starting point account for our main results.

An implicit assumption of the DDM is that evidence accumulation is triggered by the onset of the imperative stimulus, and that no integration takes place prior to this. While this assumption is reasonable for common perceptual discrimination tasks that employ supra-threshold stimuli with clearly discernible onsets (e.g. Roitman & Shadlen, 2002), it may be less valid for tasks such as ours in which detection of a low-intensity target stimulus reflects the culmination of the decision process. We acknowledge that a more appropriate conceptualization of the decision process in such contexts may be one of continuous, possibly leaky (Ossmy et al., 2013; Usher & McClelland, 2001) evidence accumulation that is triggered at trial (rather than target) onset, with temporal expectation effects arising through transient modulations of aspects of this continuous process. We propose that modelling of observed behavior on our task using the DDM nonetheless provides a highly useful simplification of such a process, for the following reasons. First, the key behavioral measures for dissociating candidate characterizations of continuous, pre-stimulus accumulation – and effects of temporal expectation thereon – are the occurrence and timing of false alarms. Yet, false alarm rates were very low in our data (Table 1) and were not affected by temporal expectation (see Results). These findings suggest that pre-stimulus accumulation was not a central determinant of decisions on our task (a point corroborated by our analysis of ERPs to expected but omitted targets; see below), was not clearly modulated by temporal expectation, and moreover that our data provided insufficient constraint for modelling this process. Second, whether one assumes target onset-triggered evidence accumulation, or continuous accumulation that is transiently modulated by temporal expectation without affecting pre-target behavior, in both cases effects of temporal expectation on non-decision time, drift rate or decision bound will have markedly different consequences for post-target detection behavior that are discernible through the DDM-based modelling scheme that we employed - specifically, a change in non-decision time primarily serves to offset the RT distribution by some constant, whereas changes in drift rate and/or bound will affect both RT and accuracy in dissociable ways. Thus, while our modelling likely provides a simplified account of how the decision process unfolds across entire trials of our task, it nonetheless permitted us to meaningfully test our key hypotheses.

**Table 1.** Trial type frequency per condition in percent.

|  | SVE | SVD | SIE | SID | LVE | LVD | LIE | LID | C |
| --- | --- | --- | --- | --- | --- | --- | --- | --- | --- |
| **Hit** | 94.55 | 91.37 | 94.17 | 90.95 | 95.33 | 93.13 | 93.81 | 91.97 | - |
| **Miss** | 4.40 | 7.47 | 4.88 | 8.96 | 2.65 | 4.76 | 3.57 | 5.71 | - |
| **False alarm** | 1.04 | 1.16 | 0.95 | 0.36 | 2.02 | 2.11 | 2.62 | 2.50 | 4.17 |
| **Correct rejection** | - | - | - | - | - | - | - | - | 95.83 |

Abbreviations: SVE: short valid easy; SVD: short valid difficult; SIE: short invalid easy; SID: short invalid difficult; LVE: long valid easy; LVD: long valid difficult; LIE: long invalid easy; LID: long invalid difficult; C: catch.

### EEG recording and preprocessing

EEG data were recorded using a BioSemi ActiveTwo system from 64 channels placed according to the international 10/20 system. Additionally, a reference electrode was placed on each mastoid, and bipolar electro-oculogram (EOG) recordings were obtained from electrodes placed approximately 1 cm lateral of the outer canthi (horizontal EOG) and from electrodes placed approximately 1 cm above and below the left eye (vertical EOG). During acquisition, impedances were kept below 30 kΩ. The EEG signal was pre-amplified at the electrode to improve the signal-to-noise ratio with a gain of 16×, and digitized at 24-bit resolution with a sampling rate of 1024 Hz. Each active electrode was measured online with respect to a common mode sense (CMS) active electrode producing a monopolar (non-differential) channel. All EEG data were analyzed in MATLAB 2012a, using the EEGLAB toolbox (Delorme & Makeig, 2004) and custom code. EEG data were re-referenced off-line to the average of the mastoid channels. Artifactual channels were interpolated with cubic spline interpolation.

In order to preserve the temporal characteristics of the slowly evolving CPP, we adopted a two-stage cleaning procedure. First, to remove drifts, the continuous EEG data were high-pass filtered offline at 0.5 Hz, and segmented from −4.8 s until +1 s surrounding target onset and surrounding response onset. An automatic algorithm detected trials that contained artifacts based on the following criteria: channel joint probability (5.5); channel kurtosis (5.5), and absolute voltage deflections (2 mV). Next, we detected eye-movement and blink artifacts using joint approximation diagonalization of eigen matrices (JADE) independent component analysis (ICA). ICA weights were subsequently stored and projected onto the raw unsegmented data to which no high-pass filter had been applied. The purpose of back projection was to prevent ICA from solely explaining variance due to large drifts in the unfiltered data and thus inaccurately identify eye-blink-related components, and to ensure that trials would not meet artifact rejection criteria due to eye blinks alone.

After removing artifactual ICA components from the unfiltered data, the data were again high-pass filtered (using a two-way least squares FIR filter with a Hanning smoothed kernel), but with a lower frequency cut-off (0.1 Hz) to preserve slow-varying components in the data. The data were segmented, and the automatic artifact detection algorithm was applied, this time with a more stringent absolute voltage deflection criterion (300 μV). All data were then manually checked for residual artifacts and such trials were removed if necessary. Following artifact rejection, the data were low-pass filtered at 6Hz. Participants for whom <50 trials remained in a single condition were excluded from further analysis to ensure that enough trials remained for single-trial analyses. This applied to 8 participants. For one participant, inadvertently no EEG data were saved to disk, making the final sample 21 participants. This exclusion criterion was determined *a priori*, and data of the excluded participants were not analyzed further. Finally, the data were re-referenced to the common average, and converted to current source density to prevent overlap between the CPP and the frontocentral negativity (Kelly & O’Connell, 2013).

### CPP identification and parameter estimation

Based on the response-locked scalp topography and following prior work (O’Connell et al., 2012; Twomey et al., 2015), all CPP-related analyses were conducted on the average of three centro-parietal EEG channels (P1, P2, and Pz). Significant deviation of the CPP from baseline was examined for each time point from 200 ms pre- to 800 ms post-target onset using permutation testing (10,000 iterations), correcting for multiple comparisons using the false discovery rate (FDR, *q* = 0.05).

To estimate CPP slope and onset, we fitted a two-part line segment that was connected by a central inflection point (Figure 2) to the data in a window starting at −200 ms pre-stimulus and ending at the time of the response (single-trial analyses) or the peak latency of the CPP (trial-average analyses). The difference in the length of the fitting window between single-trial and trial average analyses was motivated by the facts that responses should clearly demark the termination of the decision (and thus provide a good upper time limit on any single-trial electrophysiological signature of the decision process), but trial average waveforms do not have an associated response, and the peak CPP thus provided an alternate approximation of the point at which the decision process was terminated. The fitted pre-inflection line segment was constrained to have zero slope and amplitude. The two free parameters of this piece-wise function (time of inflection point and slope of post-inflection line) were fit via the Nelder-Mead Simplex optimization routine (via the *fminsearchbnd* MATLAB function), which minimized the sum of squared residuals between the fit and the observed CPP. The latency of the fitted inflection point was taken as the onset latency of the CPP, and the slope of the line segment following the inflection point was taken as the slope of the CPP. In single-trial analyses, trials with unrealistic onset latencies (prior to stimulus onset or >600 ms following stimulus onset) and trials with an unrealistic slope (zero or negative) were excluded from single-trial analysis. Although such trials were rare (average percentage of excluded trials: 0.21%; range: 0 – 0.62%), it was deemed necessary in order to avoid contamination by residual artifacts at the start of the trial and motor-related artifacts towards the end of the trial. One participant did not show a clear peak in the CPP, so for that participant we used the time of maximum first-order derivative (i.e., the time point where CPP build-up rate started to decline) as a substitute for peak latency for fitting in trial-average analyses. In cases where we used slope in between-participant correlations, we normalized the CPP amplitude across participants to counteract arbitrary differences due to scalp conduction properties, electrode impedances, and other sources of spurious variance prior to computing slope. This method of estimating CPP parameters yielded accurate fits to the data: the correlation between estimated CPP and data was 0.98 (range: 0.95 – 0.99).

**Figure 2.**
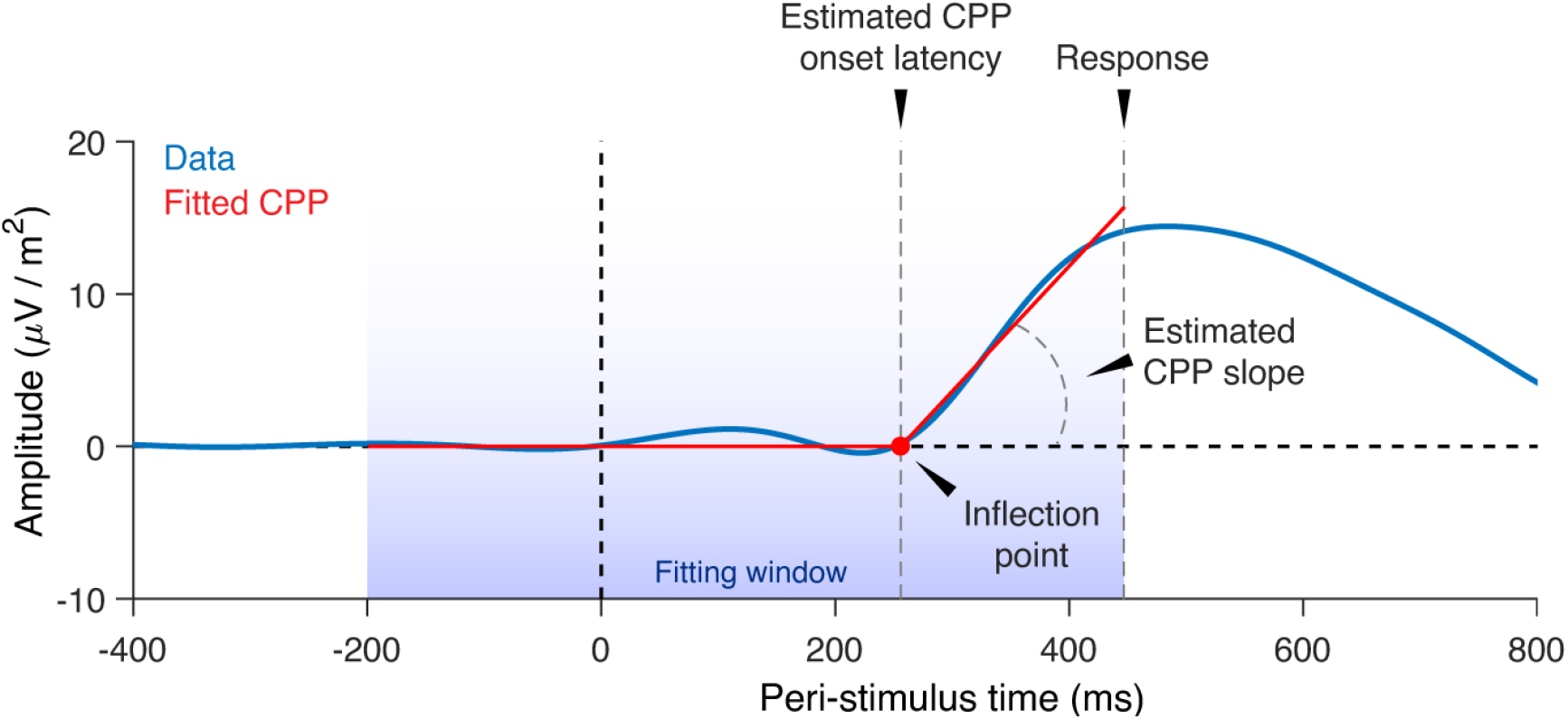
Method for estimating CPP slope and onset. Two straight lines that share an inflection point were fitted to the data with the constraint that the pre-onset part must have a slope and amplitude of zero, and the post-onset part must have a positive slope. The window for fitting was from 200 ms before stimulus onset until the time of response (for single-trial analyses) or until the peak latency of the CPP (for trial-average analyses).

Because the fitting window for parameter estimation at the single-trial level varied with RT, and RT in turn varied as a function of task condition, it was conceivable that estimated CPP onset would vary as a function of task condition by chance alone. Therefore, we computed a permuted null distribution by randomly sampling a time point within the fitting window for each trial 10,000 times. We then computed a trial-average value, separately for each condition and iteration of the permutation test, and subsequently averaged across participants. A *p* value for the true participant-average between-condition difference in estimated CPP onset was then computed by counting the number of observations in the permuted null distribution equal to or larger than the true value. Thus, the *p* value reflects the significance of the effect of condition on CPP onset latency, beyond the effect of between-condition differences in the fitting window alone. In addition to the *p* value, we report a 95% confidence interval (CI) around the mean difference between conditions, computed by summating the true mean difference between conditions and the 5^th^ and 95^th^ percentile of the permuted null distribution.

### Posterior α band power

We used Morlett wavelets to decompose the EEG data into its spectral representation. The wavelets were linearly spaced from 1 to 30 Hz with a cycle range from 3 to 12 (and were defined as in van den Brink, Wynn, & Nieuwenhuis, 2014). Power for each trial was expressed as a percentage change from the average of a −400 ms to −100 ms, frequency-specific, pre-stimulus baseline. Baseline power was calculated as the average across channels and conditions during the baseline period, in order to preserve potential between-condition differences in power prior to stimulus onset as well as their topographical distribution. We then averaged power in the α band (9-12 Hz) across 14 parieto-occipital channels, selected based on prior work on the relationship between α power and temporal expectations (Rohenkohl & Nobre, 2011).

We first examined if α power was modulated by expectations about stimulus onset by comparing valid to invalid trials with a short CTI, using nonparametric permutation testing (10,000 iterations) and expected larger α power around stimulus onset on invalid trials (i.e., when a target was presented but the participant did not expect it). We then sorted trials by α power in a −25-ms to 75-ms window surrounding target onset (i.e., the earliest window that showed an effect of expectation on α power), computed the average CPP and RT for three bins of trials of equal size, and estimated CPP slope and onset using the fitting procedure described above. We performed the binning procedure separately for each of the eight conditions of the task design to ensure that a potential relationship between α power and CPP parameters was not driven by between-condition differences in RT that covaried with α power (due to the task design), but instead reflected intrinsic trial-by-trial covariation within a given condition. Finally, for each participant we averaged across the eight conditions and then fitted a straight line to the values of RT, CPP slope and CPP onset across the three bins, and compared the distribution of slopes across participants to zero using permutation testing (10,000 iterations).

### Experimental Design and Statistical Analyses

The study followed a within-participants design with one session per participant. RT was analyzed with a repeated measures ANOVA. Error rates were analyzed with generalized mixed regression modeling (see section *Behavioral data analysis*). Effects of expectation on DDM parameters were assessed with a hierarchical fitting procedure (see section *Hierarchical drift diffusion modeling*). Effects of expectation on CCP parameters were examined using non-parametric permutation testing (see section *CPP identification and parameter estimation*). Effects of expectation on alpha power and the relationship between alpha power and CPP parameters and response time were examined using permutation testing (see section *Posterior α band power*). This study was not pre-registered.

### Data and code availability

The raw and processed data, as well as code to reproduce the results are publicly available without restriction [link inserted upon acceptance].

## RESULTS

Twenty-one individuals each performed 920 trials of a speeded, visual target-detection task in which perceptual difficulty was manipulated by adjusting the luminance of the target stimulus (Figure 1a). Temporal expectations about the timing of the target were manipulated using auditory cues. On each trial, one of two cue tones probabilistically predicted the onset time of the subsequent target. One tone was followed by an early target (1350 ms after the cue) on 80% of the trials and a late target (2700 ms after the cue) on 20% of the trials. For the other tone, these contingencies were reversed. Participants were instructed to use the temporal information signaled by the cue to optimize their RTs.

### Temporal expectations influence task performance

The behavioral results were as expected, displaying clear effects of temporal expectation and task difficulty on RT (Figure 1b). Repeated-measures ANOVAs yielded significant main effects on RT of validity (*F*(1,20) = 9.70, *p* = 0.005; 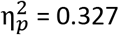), CTI (*F*(1,20) = 68.33, *p* < 0.001; 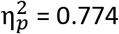), and difficulty (*F*(1,20) = 75.40, *p* < 0.001; 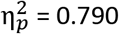). No interaction effects were significant, except the interaction effect of validity and CTI (*F*(1,20) = 32.11, *p* < 0.001; 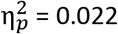). On short CTI trials, participants responded more quickly when they expected the target than when they did not expect it (*t*(20) = 6.40, *p* < 0.001; BF = 6401.65)—the typical temporal cuing effect. On long CTI trials we did not find an effect of cue validity on RT: On invalidly cued long-CTI trials the target hazard rate increases over time, despite a proportion of catch trials, leading participants to reorient their attention toward the long CTI, thus mitigating a temporal cuing effect (Correa et al., 2006). Thus, in line with previous findings (Coull & Nobre, 1998; Nobre et al., 2007), RTs for the short CTI decreased with increasing target probability, indicating that participants used the temporal information conveyed by the cues.

Results from a generalized mixed regression model predicting the proportion of misses (i.e. go trials without a response; Table 1) based on validity, difficulty and CTI and all interactions between these predictors showed significant main effects of validity (χ^2^(1) = 4.87, *p* = .027, odds-ratio = 1.18), of difficulty (χ^2^(1) = 64.94, *p* < .001, odds-ratio = 1.90), and CTI (χ^2^(1) = 31.05, *p* < .001, odds-ratio = 1.96). There were no significant interaction effects, (all *p*-values > .49). We subsequently tested whether there was an effect of validity for either of the CTIs considered separately. This was not the case, both when collapsing across difficulty levels (short CTI: χ^2^(1) = 1.60, *p* = 0.21, odds-ratio = 1.18; long CTI: χ^2^(1) = 3.75, *p* = 0.053, odds-ratio = 1.23), and for the individual difficulty levels alone (short CTI, easy: z = −0.57., *p* = 0.57, odds-ratio = 1.11; short CTI, difficult: z = −1.16, *p* = 0.25, odds-ratio = 1.18; long CTI, easy: z = −1.58., *p* = 0.12, odds-ratio = 1.41; long CTI, difficult: z = −1.22, *p* = 0.22, odds-ratio = 1.23). We next examined the proportions of false alarms (i.e. trials with a response prior to target onset; Table 1). Because for this type of error participants responded prior to stimulus onset and thus never saw the stimulus, we collapsed across difficulty levels. Results showed a main effect of CTI (χ^2^(1) = 16.07, *p* < .001, odds-ratio = 5.98), but not of validity (χ^2^(1) = 0.12, *p* = 0.73, odds-ratio = 1.26). Although the interaction between CTI and validity was significant (χ^2^(1) = 4.53, *p* = .033, odds-ratio = 0.46), the simple main effects of validity were not significant (short CTI: z = 1.70., *p* = 0.09; long CTI: z = −1.33., *p* = 0.18).

Thus, the predominant effect of temporal expectation was manifested in RT, on the short CTI, where participants responded faster when they expected the stimulus compared to when they did not expect it. The effect of cue validity on RT at the short CTI (Δ 32 ms, SD 23 ms) and the effect of difficulty (Δ 31 ms, SD 16 ms) were statistically indistinguishable (*t*(20) = 0.20, *p* = 0.84), with a Bayes factor of 0.23, indicating ‘substantial’ evidence for the null hypothesis of no difference between these effect sizes (Wetzels & Wagenmakers, 2012). This null finding enabled a fair dissociation between the distinct computational parameters that we assumed to underlie these RT effects (see below).

### Drift diffusion modelling supports decision onset account

We examined the impact of temporal expectations on latent aspects of decision-making by fitting the drift diffusion model (DDM) to participants’ behavioral data, using a hierarchical Bayesian model fitting procedure (see *Materials and Methods*; Wiecki et al. 2013). The DDM is a prominent mathematical model of simple decisions like those made in our target-detection task, and can parsimoniously account for RT distributions and choice data across a wide array of tasks (Forstmann et al., 2016). The model assumes that noisy sensory evidence is accumulated over time until one of two opposing bounds is reached, at which point a decision is made in favor of the corresponding decision. In the application of the DDM that we used (Gomez et al., 2007; Zhang et al., 2016), the upper and lower bounds correspond to a ‘go decision’ (a decision to execute a response, as expected when a target is presented) and an implicit ‘no-go decision’ (a decision to withhold an overt response, as expected when a target is not presented; Figure 3a). Previous work has found that drift diffusion models with an implicit boundary for no-go decisions provide a better fit of go/no-go task data than single-threshold variants of the model (Gomez et al., 2007). Core DDM parameters include the drift rate, *v* (which is determined by the quality of the sensory evidence and relates monotonically to the mean rate of evidence accumulation toward the correct decision bound); boundary separation, *a* (the distance between the two decision bounds, which captures response caution), and non-decision time, *T*_*er*_ (capturing the time needed for sensory encoding and response execution).

**Figure 3.**
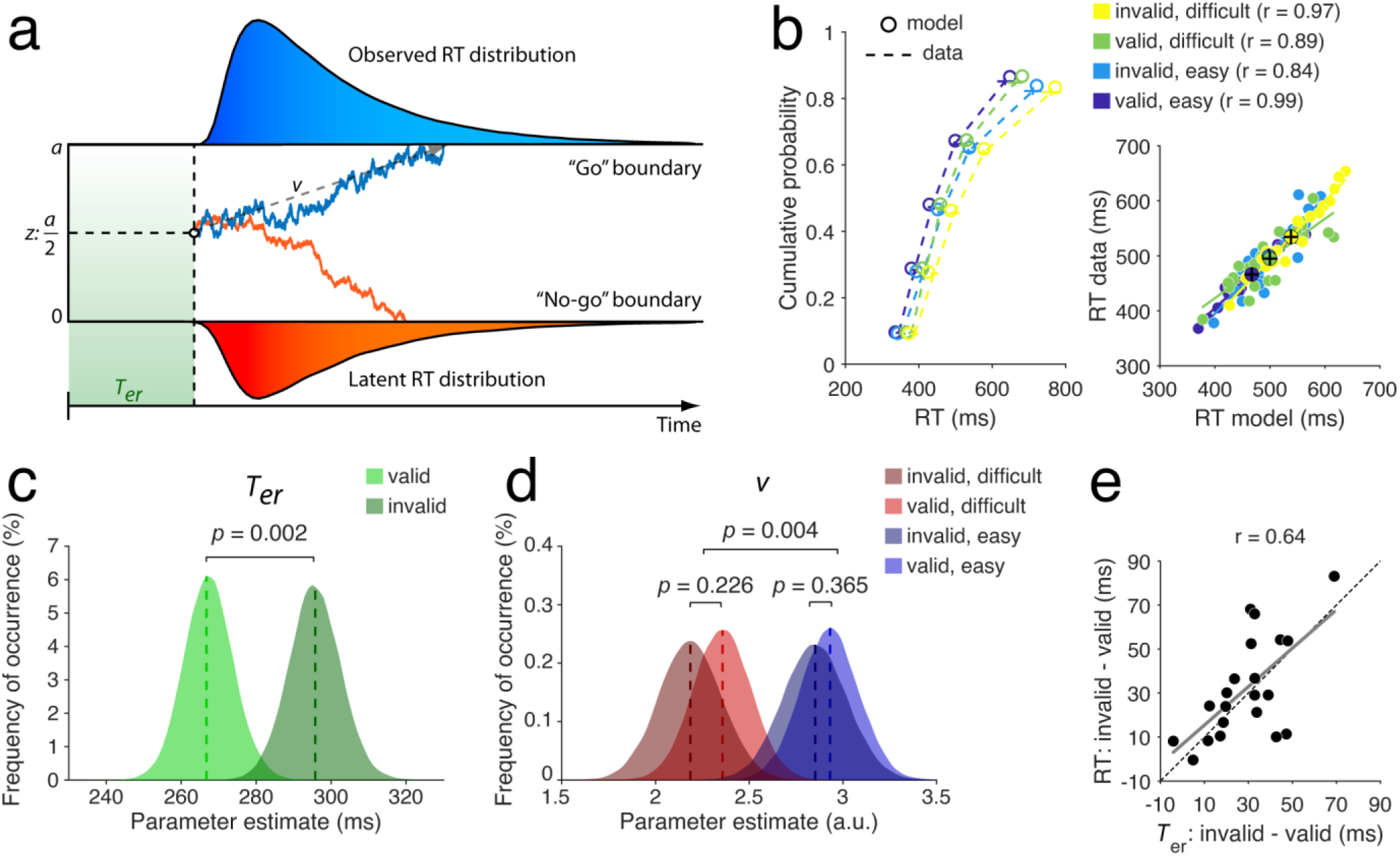
Hierarchical drift diffusion modeling results. **a)** Schematic and example traces of the go/no-go drift diffusion model. Following a period of non-decision time (*T*_*er*_), noisy evidence accumulates over time at mean drift rate (*v*) until a threshold (0 or *a*) is reached. In our implementation, starting point (*z*) was unbiased. **b)** Empirical (data) and simulated (model) defective cumulative quantile probability (left) and mean response times (right). In the right panel, small dots show individual participants. Large dots show the group means. Error bars show the within-participant SEM. **c)** Non-decision time (*T*_*er*_) differed between valid and invalid trials. **d)** Drift rate (*v*) did not differ between valid and invalid trials, but did differ between easy and difficult trials. Vertical dotted lines in c) and d) show the average of the participant-specific maximum *a-posteriori* parameter estimates. Distributions show the group posterior. Statistical significance is reflected in the overlap between distributions. **e)** Across-participant correlation between the effects of cue validity on RT and *T*_*er*_. The dotted line shows the identity line. The solid gray line shows a least-squares regression line.

To directly contrast the predictions of the decision onset account of temporal expectations with an account wherein expectations affect evidence quality (and thus accumulation rate), we examined a model fit to data from short CTI trials in which non-decision time (*T*_*er*_) and drift rate (*v*) were free to vary as a function of cue validity. Drift rate was also free to vary as a function of difficulty, in line with common findings that differences in stimulus strength are well captured by this parameter (Ratcliff, 2002). The model accurately reproduced observed RTs, with correlation coefficients between condition-specific mean RTs in data and model > 0.84, and the empirical data fell within the 95^th^ percentile credibility interval of the model (Figure 3b). Model fits showed a significant difference in *T*_*er*_ between valid and invalid trials (*p* = 0.002, Figure 3c), indicating a temporal expectation effect on non-decisional processes. In contrast, *v* did not differ between valid and invalid trials (*p* = 0.30; Figure 3d), which is inconsistent with the evidence quality account. Instead, as expected, *v* differed between easy and difficult trials (*p* = 0.004; collapsed across validity conditions).

Several features of our results showed that the effect of cue validity on RT was primarily driven by a change in *T*_*er*_. First, the effect of cue validity on *T*_*er*_ (Δ 29 ms, SD 17 ms) and the effect of cue validity on RT (Δ 32 ms, SD 23 ms) were statistically indistinguishable (BF = 0.30). Second, the participant-specific effect of cue validity on RT was positively correlated with the effect of cue validity on *T*_*er*_ (r = 0.64, *p* = 0.002; Figure 3e). Moreover, the slope of the least squares regression line that captured this relationship (0.87; 95% confidence interval determined via bootstrapping: 0.42 – 1.17) did not differ from the slope of the identity line (1.00, reflecting an idealized case where the cuing effect on RT is fully captured by *T*_*er*_). While a cross-subject correlation between the effects of difficulty on RT and on *v* was also significant (*r* = −0.63, *p* = 0.002), by contrast, there was no significant correlation between the effects of cue validity on *v* and RT (*r* = 0.32, *p* = 0.17). Together, these findings show that *T*_*er*_, not *v,* was the primary factor that drove the cue-validity effect on RT.

To further assess the effect of temporal expectations on the different model parameters we compared a range of models (including the model described above) that differed with regard to which parameter or combination of parameters was allowed to vary as a function of cue validity. In all models, *v* was free to vary as a function of difficulty. To compare the adequacy of the seven models in explaining the observed data we used the DIC metric, a statistical criterion for model selection that takes into account model complexity. As can be seen in Table 2, the model in which only *T*_*er*_ was free to vary between valid and invalid trials provided the best fit of the data—a significantly better fit than the model ranked second (*T*_*er*_ + *a* + *v*, ΔDIC = 10) and the *v*-only model (ΔDIC = 179). Notably, the four best fitting models were all of the models in which *T*_*er*_ was allowed to vary between valid and invalid trials. Thus, our DDM analyses strongly supported an account whereby temporal expectations selectively act to hasten non-decisional processes, but do not affect the process of evidence accumulation itself.

**Table 2.**
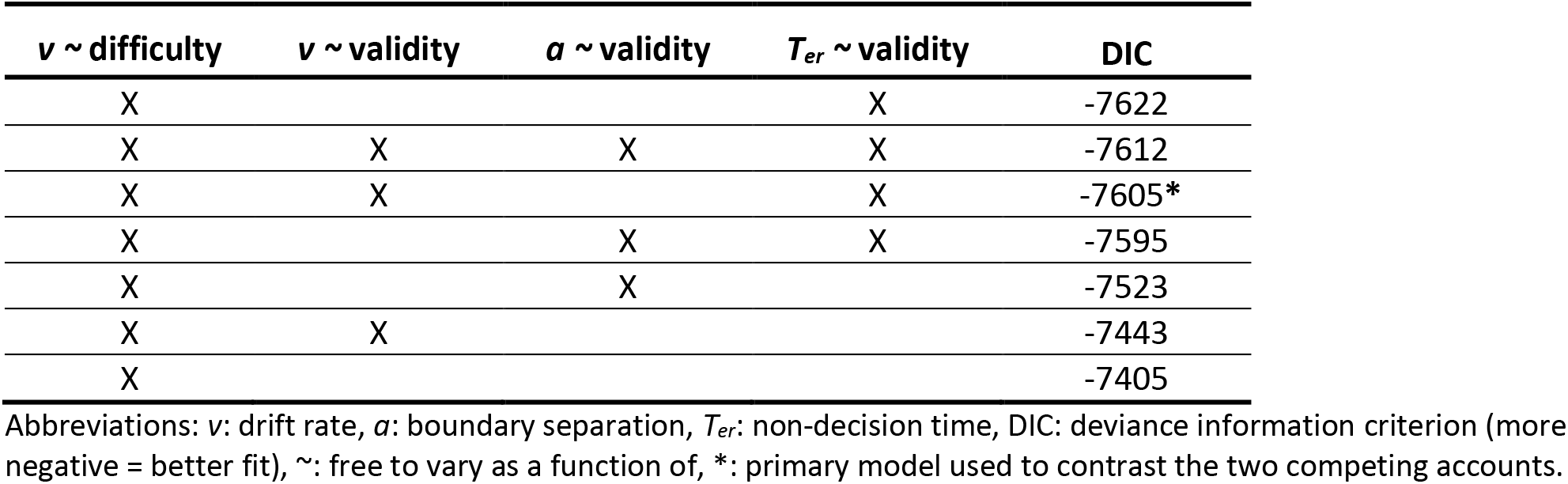
Models ranked in terms of deviance information criterion.

| $v \sim \text{difficulty}$ | $v \sim \text{validity}$ | $a \sim \text{validity}$ | $T_{er} \sim \text{validity}$ | DIC |
| --- | --- | --- | --- | --- |
| X |  |  | X | -7622 |
| X | X | X | X | -7612 |
| X | X |  | X | -7605* |
| X |  | X | X | -7595 |
| X |  | X |  | -7523 |
| X | X |  |  | -7443 |
| X |  |  |  | -7405 |

Abbreviations: $v$ : drift rate, $a$ : boundary separation, $T_{er}$ : non-decision time, DIC: deviance information criterion (more negative = better fit), ∼ : free to vary as a function of, \*: primary model used to contrast the two competing accounts.

The *v* account predicts that a change in temporal expectations should result in a concomitant change in accuracy, which we did not reliably observe at the short CTI. However, accuracy on our task was high overall (by design), which may have absorbed a potential effect on accuracy. Alternatively, it is possible that a change in *v* due to expectation occurred simultaneously with a change in *a*, leaving accuracy levels unaffected while improving RT. To exclude this possibility, we fit an additional model in which *v* and *a* were free to vary as a function of cue validity, *v* was additionally free to vary as a function of difficulty, and *T*_*er*_ was not free to vary. The DIC score of this model (−7497) was significantly higher (indicating a poorer fit) than that of the best-performing model (Table 2) in which the effects of temporal expectation were captured by *T*_*er*_ only (ΔDIC = 125). Moreover, a model in which all three parameters were free to vary as a function of validity did not yield a significant effect of validity on *v* or on *a* (*v*: *p* = 0.74, *a*: *p* = 0.23). Thus, although changes in *v* and *a* could jointly produce reduced RTs without changes in accuracy, our results indicate that the effects of temporal expectations on performance were better captured by a change in *T*_*er*_ only.

### Target detection is associated with an electrophysiological signature of decision making

Previous work has identified an electrophysiological signature of evidence accumulation during decision making tasks: a supramodal, centroparietal positivity (CPP) in the event-related potential that exhibits the typical dynamics of a build-to-threshold decision signal (Kelly & O’Connell, 2013; O’Connell et al., 2012; Twomey et al., 2015). The CPP is motor-independent: it is robustly observed in cases where no motor response is required (O’Connell et al. 2012) and its dynamics are clearly dissociable from motor signals (Twomey et al., 2015), indicating that this signal reflects decision formation and not merely motor preparation or execution. The CPP thus provided us with a neural measure by which to examine the effect of temporal expectation on decision formation (Loughnane et al., 2016).

In line with previous work, the CPP in our data showed a significant (*p* < 0.05, FDR-corrected), gradual build-up following target onset, peaked around the time of the response, and was maximal over centroparietal electrodes (Figure 4a). Moreover, the CPP was uncontaminated by visual-evoked potentials, as our task was designed to minimize sensory transients. The CPP also exhibited the typical relationship with RT that might be expected from a build-to-threshold decision variable signal. Specifically, we split each participant’s RT distribution into equal-sized fast, medium and slow bins (within CTI, difficulty, and validity conditions) and plotted the average waveforms aligned to target presentation and response execution for each bin (Figure 4b). Consistent with previous observations (Kelly & O’Connell, 2013; O’Connell et al., 2012) the onset and peak latencies of the stimulus-locked CPP directly scaled with RT, while its slope (i.e., build-up rate) was inversely proportional to RT (see Figure 5 for fits of individual participants, and cross-participant correlations with behavior and model parameters). Furthermore, the response-locked CPP reached a stereotyped amplitude at the time of response execution, consistent with fixed decision bounds. Combined, these findings are consistent with the notion that during a detection task with high accuracy, the CPP tracked the gradual accumulation of stationary sensory evidence towards a decision bound.

**Figure 4.**
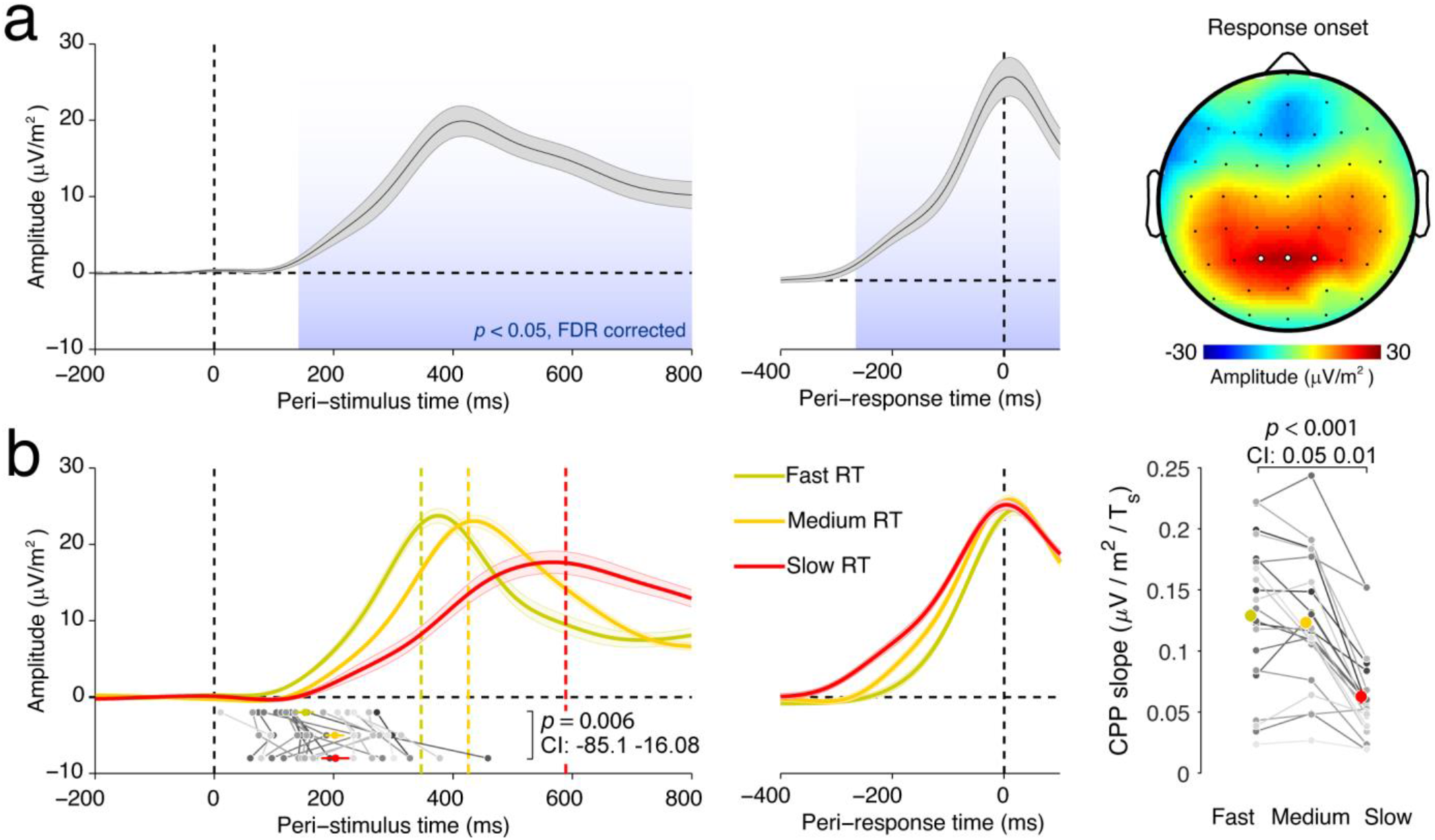
Condition-average CPP. **a)** Stimulus- and response-locked CPP waveforms and scalp topography. Error bars show the within-participant SEM. The highlighted electrodes, P1, P2 and Pz, were used for all analyses. **b)** CPP characteristics scale with response time (RT). Single trials were sorted as a function of RT and divided into equal-sized fast, medium and slow bins, separately for each participant and task condition, then averaged. Colored vertical dotted lines show the average RT per bin. Dots below the traces show the CPP onset latency. Error bars show the within-participant SEM. Gray lines show data of individual participants. CI: Permutation-derived 95% confidence interval around the mean difference between conditions.

**Figure 5.**
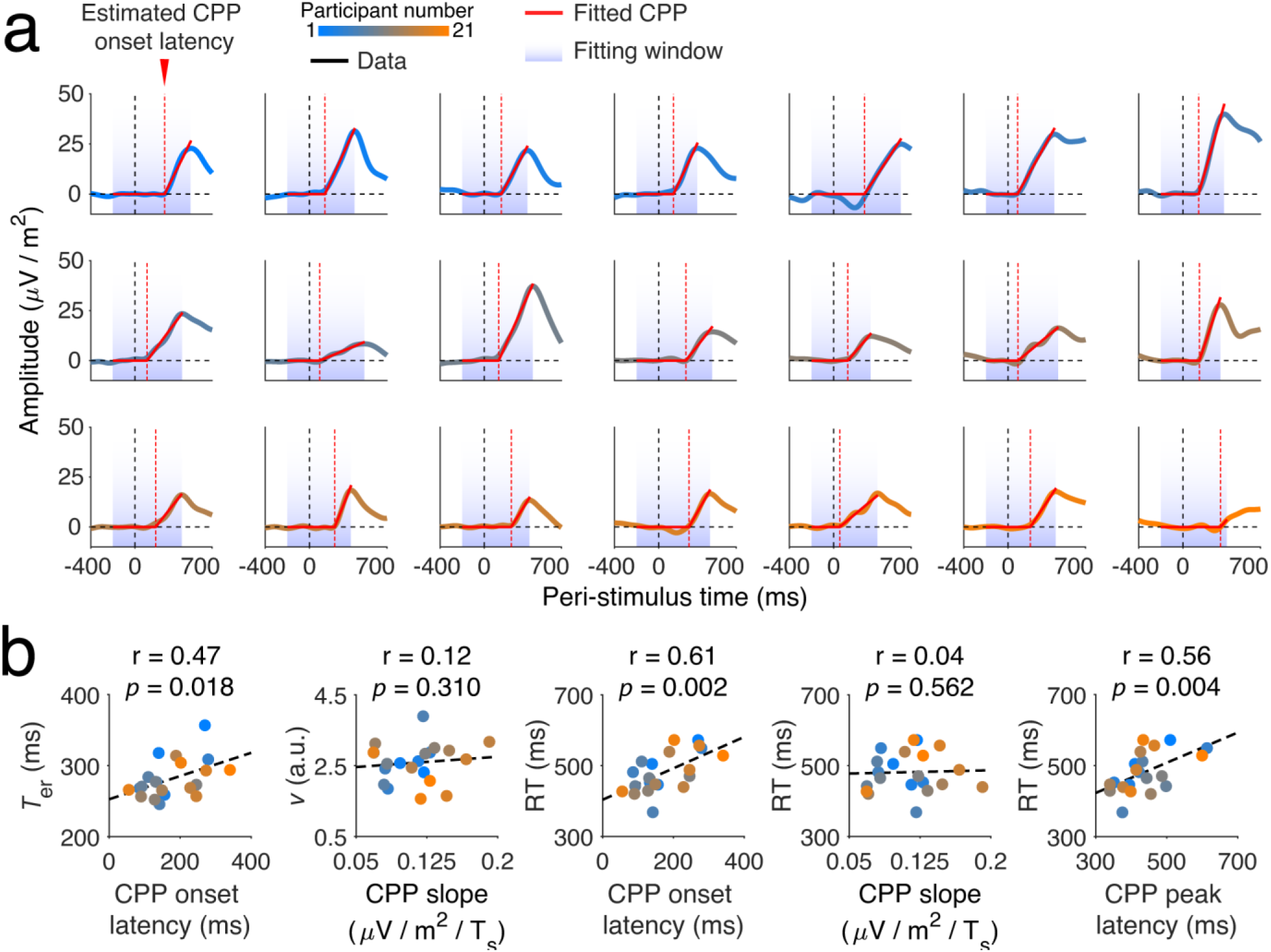
CPP fits per participant, and relationship with other measures. **a)** Condition-averaged CPP for each participant. Solid red lines show the fitted CPP traces. Red dashed lines show the estimated CPP onset latency. **b)** Cross-participant correlations between model parameters, response time (RT), and three CPP measures: CPP onset latency, CPP slope, and CPP peak latency. Prior to estimating slope, the individual CPP curves were scaled to fixed height to correct for arbitrary amplitude differences between participants.

### Temporal expectation and task difficulty have dissociable effects on CPP onset latency and slope

The decision onset account predicts that temporal expectation reduces the onset latency of the CPP. In contrast, an effect of temporal expectation on evidence quality predicts a steeper slope (i.e., build-up rate) of the CPP (Kelly & O’Connell, 2013). Trial-average analyses (Figure 6a, b) provided preliminary support for the decision onset account. Cue validity showed a trend-level effect on onset latency in the expected direction (*p* = 0.097, 95% CI: −56.71 6.23), but no effect on slope (*p* = 0.555, 95% CI: −0.01 0.01). In contrast, difficulty affected the CPP slope (*p* = 0.010, 95% CI: 0.002 0.013), reflecting the expected shallower CPP slope on difficult trials, but not the onset latency (*p* = 0.775, 95% CI: −0.23.65 7.18). This also indicated that the absence of an effect of cue validity on CPP slope was not due to a lack of sensitivity in accurately measuring CPP slope, because task difficulty and cue validity had a comparable effect on RT. Furthermore, the latency between the peak of the response-aligned CPP and the motor response (a proxy for the time between decision and response execution) did not differ between valid and invalid trials (*p* = 0.31, BF = 0.38). This suggested that the effect of cue validity on *T*_*er*_ (see previous section) was unlikely to be due to between-condition differences in post-decisional processing, and instead likely reflected differences in decision onset.

**Figure 6.**
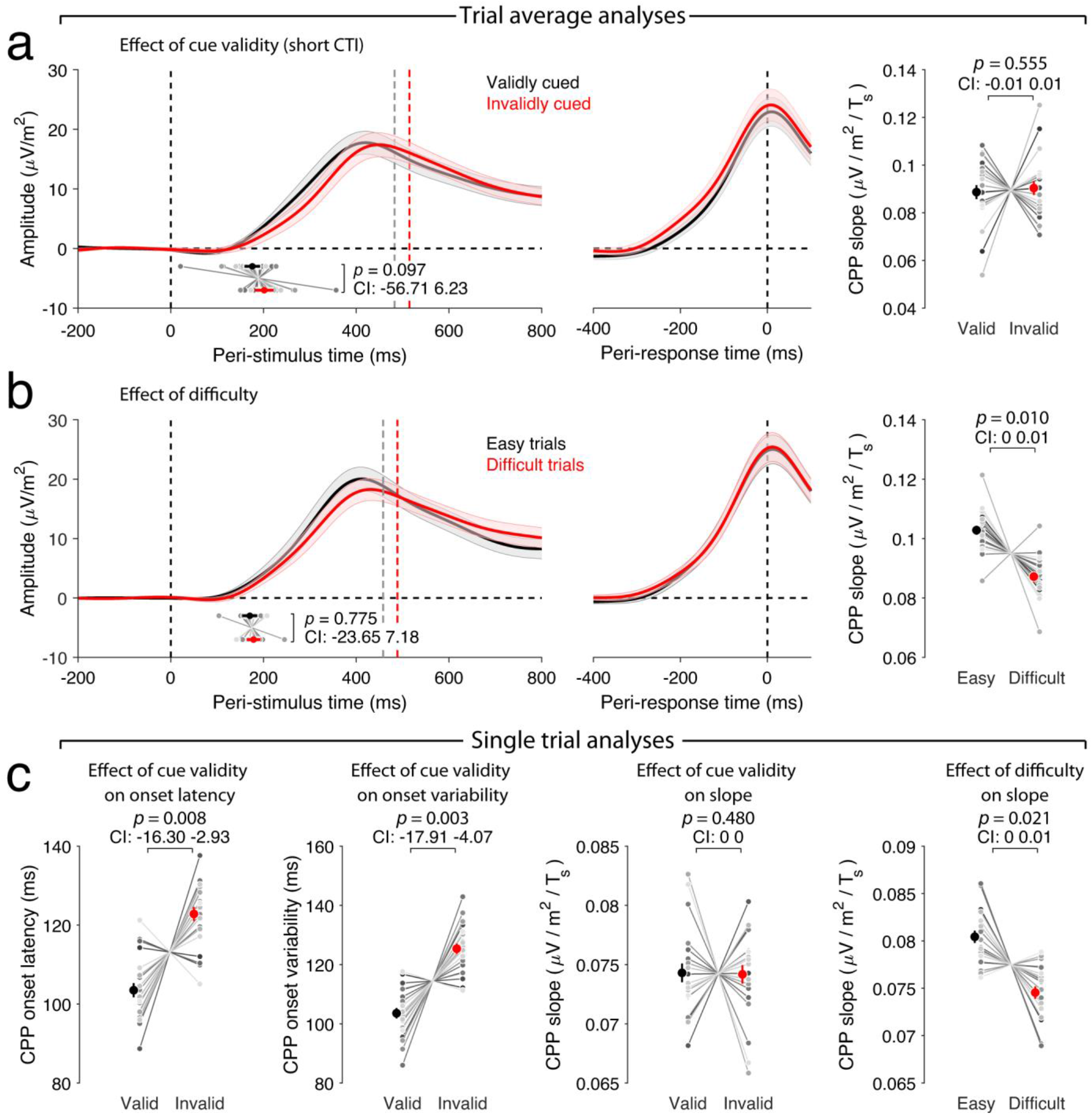
Effects of cue validity and difficulty on the CPP. **a)** Effect of cue validity, and **b)** difficulty on the trial-averaged CPP. Dots and lines below the ERP waveforms show the average estimated CPP onset latency and within-participant SEM. **c)** Comparison between CPP measures extracted at the single-trial level. Error bars reflect the within-participant SEM. Gray lines show data of individual participants, centered with respect to the condition and participant average. CI: Permutation-derived 95% confidence interval around the mean difference between conditions.

The trial-average analyses may have underestimated the true effect of validity on CPP onset latency if invalid short-CTI trials (in which the early target was not expected) were associated with larger trial-by-trial variability in onset latency. In that case, the onset latency distribution for invalid trials might have a wider left tail, which would have led to an earlier onset of the trial-averaged CPP, diminishing the difference in CPP onset latency between valid and invalid trials. To examine this possibility, we computed single-trial measures of CPP onset latency and slope (see *Materials and Methods*). Although the mean single-trial estimated CPP onset latencies were significantly correlated with trial average estimates across participants (r = 0.73, p < 0.001), these data confirmed that invalidly cued short-CTI trials were indeed associated with increased variability in onset latency compared to validly cued short-CTI trials (*p* = 0.003, 95% CI: −17.91 −4.07, Figure 6c). This was not simply a consequence of variation in the quality of CPP model fits, as the correlation between estimated CPP and data at the single-trial level did not differ between valid and invalid conditions (valid: 0.82; invalid: 0.83; *t*(20) = 0.63, *p* = 0.53, BF = 0.27). Our key predictions might therefore be addressed more reliably using single-trial analyses.

To avoid spurious effects of trial-to-trial variability on the shape of the trial-average CPP waveform, we examined the effects of temporal expectation and task difficulty on single-trial measures of CPP onset latency and CPP slope (Figure 6c). These analyses revealed that the CPP onset latency was significantly shorter on validly cued trials than on invalidly cued trials (*p* = 0.008, 95% CI: −16.30 −2.93), while the CPP slope did not differ (*p* = 0.480, 95% CI: −0.003 0.003, BF = 0.23). As expected, and in line with the trial-average analyses, CPP slope was shallower for difficult trials than for easy trials (*p* = 0.021, 95% CI: 0.0005 0.0054).

Our finding of a cue-validity effect on CPP onset latency is consistent with the notion that temporal expectation hastens decision onset. However, an alternative possibility is that on validly cued (versus invalidly cued) short-CTI trials, when the target was expected to appear after the short CTI, participants engaged in premature sampling (of noise) on a proportion of the trials (Jepma, Wagenmakers, & Nieuwenhuis, 2012; Laming, 1979; Devine et al., 2019). That is, participants may anticipate the arrival of a target stimulus and speed up responses by starting to sample information from the display at the moment when they think the stimulus will be presented. On a proportion of the trials they may start sampling too early, and if these premature sampling trials are included in the analysis, the average single-trial CPP onset latency may give the impression of speeded decision onset.

Premature sampling should lead to faster but less accurate responses, because participants will start with sampling noise. Accordingly, we discouraged premature sampling by instructing participants to avoid response errors and aborting trials on which responses were made before target onset. Anticipations were not more common on short-CTI trials when a target was expected, compared to when a target was not expected (*t*(20) = 1.39, *p* = 0.18, BF = 0.52) suggesting that any premature sampling that participants may have engaged in was not modulated by temporal expectation. To further rule out a role for premature sampling, we carried out a control analysis, focusing on the ERP waveform associated with invalidly cued long-CTI trials. On these trials participants expected the target to occur after the short CTI, but it was not presented. Given that the CPP appears to reflect accumulated evidence regardless of sign (O’Connell et al., 2012), premature sampling (relative to the true target onset latency) should result in a positive “pre-target” CPP deflection that cannot possibly reflect evoked activity, because the target is not yet presented at that time. However, our control analysis did not yield any support for this prediction (Figure 7).

**Figure 7.**
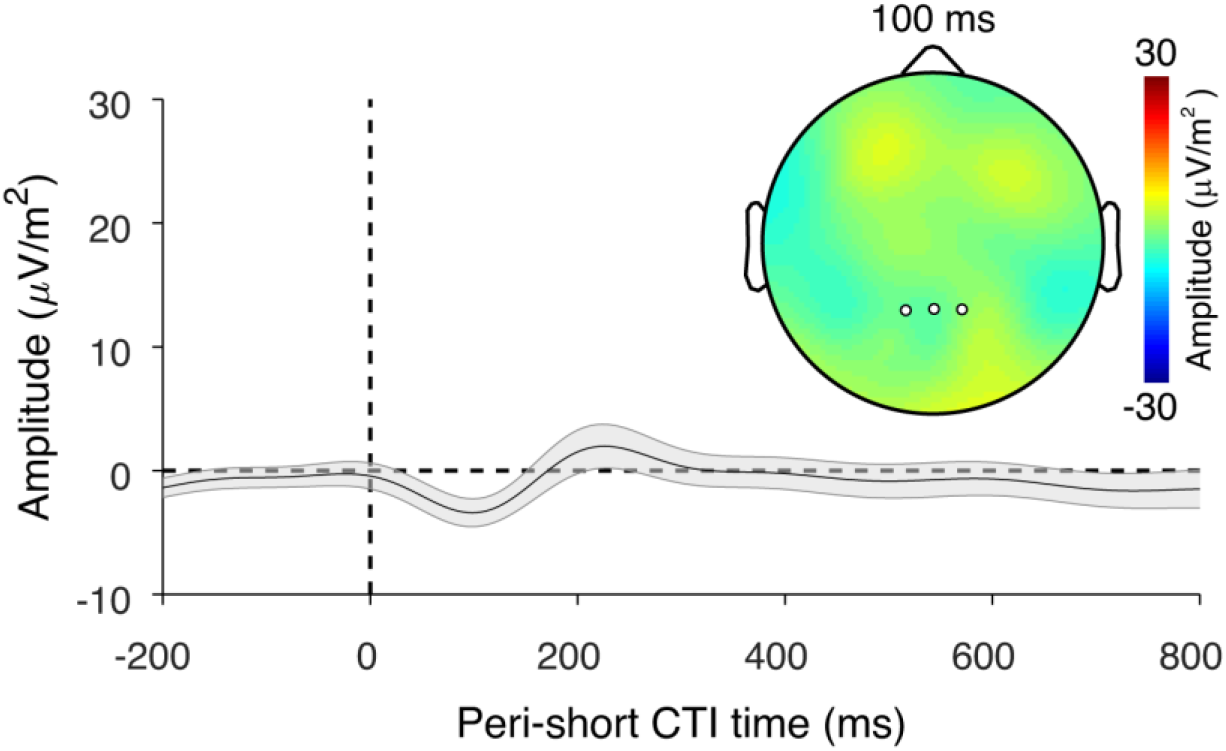
Control analysis to rule out premature sampling. The figure shows the ERP for invalid long-CTI trials, at the time points surrounding the short CTI (i.e., when a target was expected, but not presented), relative to a 200 ms pre-cue baseline. If participants were starting to accumulate evidence prior to stimulus onset, we would expect to see a positive deflection around stimulus onset, but no time points were significantly different from zero (all FDR *p* > 0.05). Color and axis limits are as in Figure 4a. The error bar shows the within-participant SEM.

### CPP onset latency covaries with behaviorally relevant fluctuations in peri-stimulus α power

In search of a mechanism underlying the effect of temporal expectations on decision onset, we examined a potential endogenous determinant of CPP onset latency that might account for trial-to-trial variability in CPP onset latency and corresponding fluctuations in task performance. In particular, we used EEG time-frequency analysis to examine the power of ongoing neural oscillations in the α band (9-12 Hz) over posterior cortical areas, reflecting rhythmic fluctuations in cortical excitability (Jensen & Mazaheri, 2010; Pfurtscheller, 2001). A large number of studies have found that the amplitude of α-band oscillations around the onset time of a target stimulus predicts subsequent task performance (Ergenoglu et al., 2004; Mathewson, Gratton, Fabiani, Beck, & Ro, 2009; Thut et al., 2006; van Dijk, Schoffelen, Oostenveld, & Jensen, 2008), in line with the well-accepted notion that the neural and behavioral responses to a stimulus depend on the cortical state at the time of stimulus presentation. Importantly, the power and phase of α-band oscillations around the time of a stimulus can be modulated by top-down attention to optimize the processing of that stimulus (Samaha, Bauer, Cimaroli, & Postle, 2015; van Diepen, Cohen, Denys, & Mazaheri, 2015).

The top-down modulation of the power of oscillatory α-band activity has been proposed to play a pivotal role in performance on temporal expectation tasks. Previous EEG research has found that α power is reduced just prior to, or at, the expected onset time of a target event (Breska & Deouell, 2017; Praamstra, Kourtis, Kwok, & Oostenveld, 2006; Rohenkohl & Nobre, 2011), yielding enhanced cortical excitability over the time interval in which the target event is expected. This signature of temporally focused attention, often referred to as α desynchronization, was also present in our data. Power in the α band was maximal over posterior scalp regions (Figure 8a) and specifically posterior α was modulated by expectations about stimulus onset (Figure 8b), with lower α power preceding the onset of an expected stimulus compared to an unexpected stimulus, possibly reflecting stronger anticipatory α desynchronization. This led us to ask whether fluctuations in this signal were associated with fluctuations in CPP onset latency, our neural marker of decision onset. To address this question, we binned trials based on single-trial, peri-stimulus posterior α power within each cell of the task design, and then averaged across cells to make sure that any difference between bins in α power was unrelated to task manipulations, including the cue validity manipulation, but instead reflected endogenous trial-by-trial variability.

**Figure 8.**
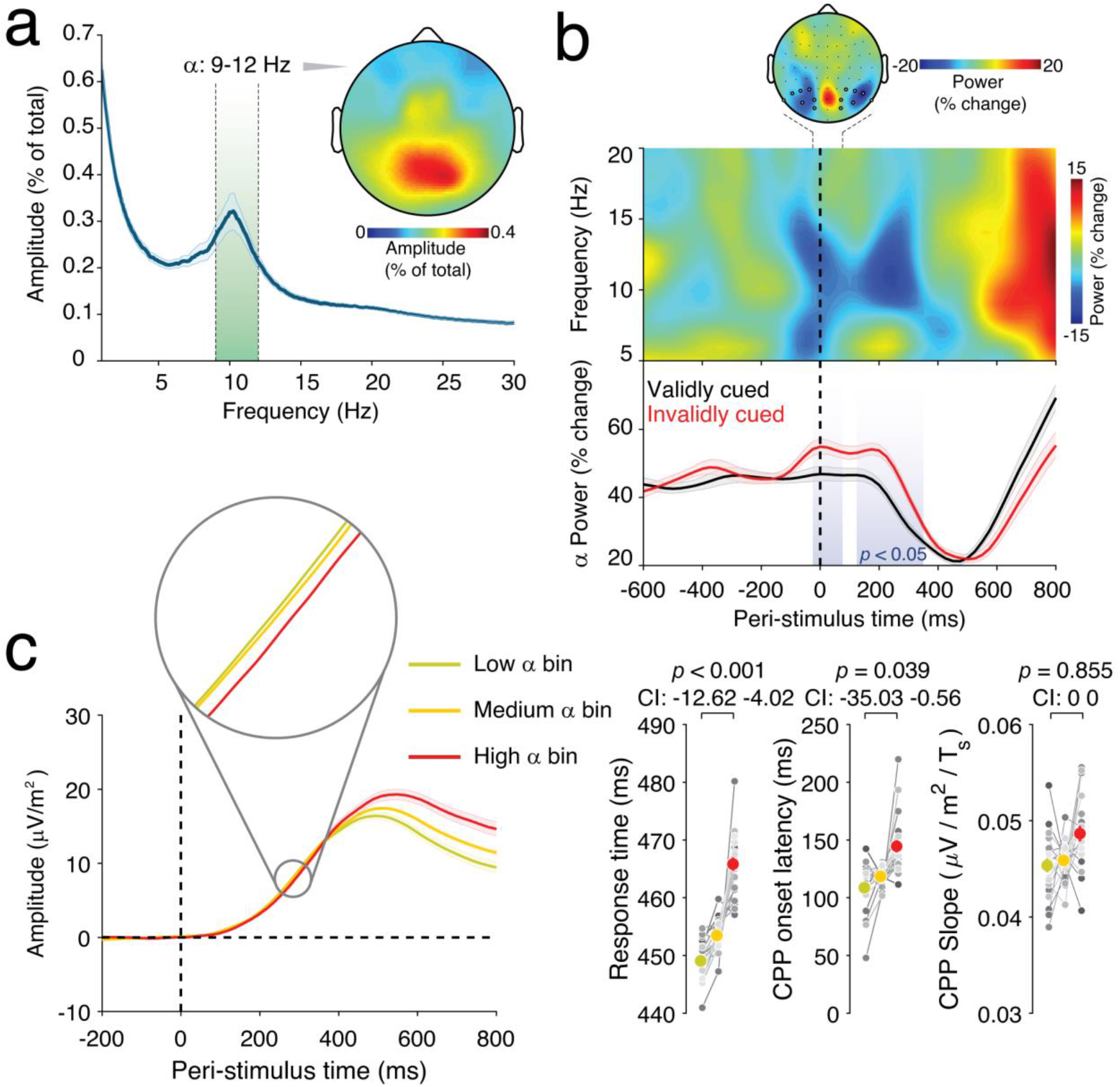
Relationship between posterior α (9-12 Hz) power, temporal expectations and CPP parameters. **a)** Power spectrum and topography of α power. **b)** Comparison of power across time and frequency for valid versus invalid trials. Blue colors indicate lower power for valid trials. The topographical plot shows the comparison of power in the α (9-12 Hz) band between valid trials and invalid trials on the short CTI, collapsed across difficulty levels; highlighted channels are those averaged over to create remaining panels, chosen independently, based on Rohenkohl and Nobre (2011). Shaded areas in the lower panel indicate a significant (p < 0.05, uncorrected) difference between valid and invalid trials). **c)** Trial- and condition-averaged CPP binned by peri-stimulus α power (*left*), and effect of α bins on RT, CPP onset latency and CPP slope (*right*). Error bars show the within-subject SEM. Gray lines show data of individual participants, centered with respect to the condition and participant average. CI: Permutation-derived 95% confidence interval around the mean difference between conditions.

A statistical comparison between bins indicated that the differences in peri-stimulus posterior α power were behaviorally relevant, as decreases in α power were associated with faster RTs (Figure 8c; *p* < 0.001, 95% CI: −12.62 −4.02). Importantly, decreased α power was also associated with a reduced CPP onset latency (*p* = 0.039, 95% CI: −35.03 −0.56), but not with differences in CPP slope (*p* = 0.855, 95% CI: −0.004 0.0008) (Figure 8c). Identical results in terms of direction and significance were found using an alternate frequency band (9-14 Hz). As expected, long CTI trials, which showed no significant effect of cue validity on RT, also showed no significant effect of cue validity on α power. Combined, our results indicate that top-down modulation of α-band activity putatively modulates cortical excitability in a way that expedites the onset of evidence accumulation of expected stimuli.

## DISCUSSION

Temporal contingencies in sensory input, such as music, speech and other temporal sequences, provide a critical source for temporal predictions formed in the brain, considerably improving the speed and accuracy of responding to unfolding events. Here, we used mathematical modelling of behavior and neural signal measurements to identify how temporal expectations enhance perception.

All model-based analyses suggested that the cue-validity effect, our behavioral measure of temporal expectations, was specifically driven by changes in non-decisional processes (*T*_*er*_), and not by possible effects on the decision process (including an effect on evidence quality and, accordingly, the mean rate of accumulation, *v*). Specifically, model fits as well as model-selection procedures supported the *T*_*er*_ account. Furthermore, the cue-validity effects on *T*_*er*_ and RT were highly correlated and almost identical in size. No such support was obtained for an account whereby temporal expectations enhance the quality of the sensory evidence (Rohenkohl et al., 2012a). Although non-decision time includes the time needed for both sensory encoding and response execution, previous studies have found that temporal expectation has negligible effects on response execution time, as indicated by the interval between the onset of the lateralized readiness potential and the overt response (Hackley, Schankin, Wohlschlaeger, & Wascher, 2007; Müller-Gethmann, Ulrich, & Rinkenauer, 2003; but see Tandonnet, Burle, Vidal, & Hasbroucq, 2006). Likewise, we found that the time difference between the peak latency of the CPP and the motor response, a proxy for the duration of response execution, did not differ between valid and invalid trials. Although this metric is not without limitations because the CPP may in some cases continue to accumulate post-decision (Steinemann et al., 2018), these results indicate that our temporal expectation effect on *T*_*er*_ primarily reflects pre-decisional rather than post-decisional processing time.

Our study is the first to complement mathematical modelling with detailed EEG analyses regarding the effects of temporal expectation on decisional and non-decisional process. Such neural evidence is critical, given reported discrepancies between results from sequential-sampling models and neural signatures of decision formation (McGovern et al., 2018; Spieser et al., 2018). These additional analyses of the CPP allowed us to examine the effect of temporal expectation on the onset and rate of decision formation in the brain (Loughnane et al., 2016), and yielded further support for the decision onset account. Single-trial analyses, necessary to avoid a measurement artifact in the trial-average CPP, revealed a shortened CPP onset latency on validly cued trials, consistent with the notion that temporal expectation hastens decision onset. In contrast, valid and invalid trials did not differ in CPP slope, suggesting that temporal expectation did not affect the quality of the sensory evidence or other processes influencing the rate of decision formation. Control analyses excluded premature sampling of the stimulus array prior to target onset (Jepma et al., 2012; Laming, 1979; Devine et al., 2019) as an alternative explanation of these results.

EEG α-band power is thought to reflect local cortical excitability (Jensen & Mazaheri, 2010; Pfurtscheller, 2001) and has previously been shown to track temporal expectations, enhancing perception of events occurring at expected moments (Heideman et al., 2018; Rohenkohl & Nobre, 2011; Zanto et al., 2011). In our study, peri-stimulus posterior α power was also behaviorally relevant, showing a strong positive relationship with RTs. The novel insight provided by our data was that peri-stimulus α power also covaried with CPP onset latency (and not with CPP slope), suggesting that peri-stimulus posterior α power is a key determinant of decision onset, presumably contributing to the perceptual facilitation associated with temporal expectation. This facilitation may be brought about by top-down signals that modulate visual cortex activity in order to expedite target processing, and result in an earlier start of the decision process (and therefore a shorter CPP onset latency). Notably, α is the dominant frequency band for feedback signaling from higher-order cortical regions to lower-level visual cortex (Michalareas et al., 2016; van Kerkoerle et al., 2014). Thus, it is possible that α power is a signature of a top-down modulatory process: co-variation between posterior α power and CPP onset latency may reflect fluctuations in top-down signaling in accordance with expectations about the appearance of a target.

An important open question is how temporal expectation hastens decision onset. An implicit assumption of many sequential sampling models is that evidence accumulation begins automatically, immediately when sensory encoding is completed. Under this assumption, an expectation-induced reduction of decision onset latency necessarily stems from faster sensory encoding. However, recent literature suggests that decision onset may be de-coupled from sensory encoding, and possibly under strategic control (Teichert, Grinband, & Ferrera, 2016). Hence, it is possible that our findings reflect a strategic reduction in decision onset that is unrelated to sensory encoding time. While the CPP allowed us to directly estimate the onset of decision formation, it did not allow us to directly measure the process of sensory encoding. Thus, future efforts to combine measures of both decision onset and sensory encoding may prove fruitful in answering the question of whether temporal expectation hastens decision onset through faster sensory encoding, or through strategic control of the onset of the accumulation process.

There are prominent discrepancies between the current results and previously published model-based analyses of behavioral temporal expectation effects (Cravo et al. 2013; Rohenkohl et al., 2012a; Vangkilde et al., 2012). Vangkilde and colleagues (2012; Vangkilde, Petersen, & Bundesen, 2013) found that temporal expectations enhanced (unspeeded) visual letter discrimination and analyzed these behavioral data using the *theory of visual attention* (TVA). These analyses suggested that the improvement in perceptual discrimination was caused by an increased rate of evidence accumulation (TVA parameter: perceptual processing speed), and not by faster sensory encoding (TVA parameter: temporal threshold for conscious perception). Likewise, Nobre and colleagues found that temporal expectations enhanced visual contrast sensitivity, and reported fits of a sequential-sampling model suggesting that this enhancement in perceptual discrimination was caused by an increased quality of the sensory evidence (Cravo et al. 2013; Rohenkohl et al., 2012a).

Nobre and colleagues (Cravo et al. 2013; Rohenkohl et al., 2012a) modeled their behavioral data using a diffusion model that was developed to simultaneously fit psychometric (accuracy) and chronometric (RT) functions (Palmer, Huk, & Shadlen, 2005). However, we argue that this diffusion model was not entirely appropriate with regard to the task performed by the participants. Specifically, in both studies participants were asked to discriminate the orientation of visual gratings (Gabor patches) at seven contrast levels, resulting in accuracy levels spanning from near chance to near perfect. Each trial consisted of a stream of noise patches and infrequent visual gratings (targets) presented either with a fixed, rhythmic (high temporal expectation) or jittered, arrhythmic (low temporal expectation) stimulus onset asynchrony. Critically, each target was presented for only 50 ms and then followed by a noise patch after 350 ms. This means that the task potentially violated an assumption underlying the diffusion model of Palmer and colleagues, namely that the evidence on the screen does not change during the decision process. The consequences of the violation of this stationarity assumption are unclear. The brief target duration and the short time until the next noise patch meant that on that task, there was very limited time for participants to secure a robust visual short-term memory trace of the decision-relevant stimulus feature, which in turn provides input to the decision process once the stimulus is no longer visible (cf. Ratcliff, Smith, Brown, & McKoon, 2016; Rohenkohl et al., 2012b; Smith & Ratcliff, 2009). In such conditions, expedited non-decision time could well be beneficial for discrimination accuracy in that it would allow more time for the creation of a high-quality memory trace before the perturbance of the subsequent noise patch (cf. Nieuwenhuis, Jepma, & Wagenmakers, 2012; Rolke & Hofmann, 2007). Importantly, such a change in accuracy is often diagnostic of a change in drift rate, while generally inconsistent with a change in non-decision time. Thus, a potentially violated assumption may have led the model to misattribute an effect on non-decision time to a change in drift rate. Taken together, these arguments invite caution when interpreting previous modelling results (Cravo et al. 2013; Rohenkohl et al., 2012a) in terms that are explicit about the locus of the effect of temporal expectation on performance.

The discrepancy between our results and those of Vangkilde and colleagues (2012) also deserves further theoretical consideration. To our knowledge, the formal relationship between diffusion models and the TVA, including the mapping between their model parameters, has not been studied yet. Given the model assumption and mimicry issues discussed here, it is critical that we obtained solid neural evidence for the decision onset account.

Another plausible explanation for the discrepancy between the current results and previous work concerns differences between studies in the structure of temporal information provided to participants. Nobre and van Ede (2017) distinguish four types of informative temporal structures that are commonly found in the environment and manipulated in experimental tasks: (i) temporal associations—that is, predictive temporal relations between successive stimuli, such as the auditory cue and the visual target in our study; (ii) hazard rates (Vangkilde et al., 2013); (iii) rhythms; and (iv) sequences—recurring temporal structures that are more complex than simple rhythms. Although there is some evidence that temporal expectations based on these structures rely on similar brain mechanisms (Correa & Nobre, 2008), the differences and similarities between the types of temporal structures and associated brain mechanisms are only beginning to be investigated (Bouwer, Honing, & Slagter, in press; Breska & Deouell, 2017; Nobre & van Ede, 2017; Shalev, Nobre, & van Ede, 2019). Interestingly, the manipulations of temporal expectation of Vangkilde and colleagues (2012, 2013) and Nobre and colleagues (Cravo et al. 2013; Rohenkohl et al., 2012a) were based on hazard rates and rhythms, respectively. In contrast, behavioral studies that used temporal associations (i.e., temporal cuing or a foreperiod manipulation) have reported evidence that temporal expectations specifically affect decision onset and not the decision process itself (Bausenhart et al., 2010; Seibold et al., 2011; Jepma et al., 2012). Future research should examine the possibility that different types of temporal structures call upon dissociable temporal expectation mechanisms. Additionally, it remains to be investigated to what extent our findings generalize to perception in other sensory modalities (Ball et al., 2017; Lee et al., 2018; Herbst & Obleser, 2019).

Our findings contribute to a growing literature documenting the usefulness of the CPP as a neural marker of perceptual decision formation (O’Connell et al., 2018), but there are also important outstanding questions regarding the methodological limitations and functional significance of the CPP (Urai & Pfeffer, 2014). Furthermore, recent work has put forward the peak latency of the N200 component as an alternative EEG marker of the onset of the evidence accumulation process (Nunez, Gosai, Vandekerckhove, & Srinivasan, 2019). Additionally, although our study focused on perception, there is marked evidence that temporal expectation can also affect aspects of motor processing (e.g., Fecteau & Munoz, 2007; Hackley & Valle-Inclán, 2003; Nobre et al., 2007), albeit perhaps not the *duration* of these processes. Lastly, future research may shed more light on the precise mechanistic origin of trial-by-trial co-fluctuations in posterior α power around the time of stimulus presentation and the onset of decision formation.

## ACKNOWLEDGEMENTS

We thank Konstantinos Tsetsos for helpful comments on an earlier version of this article. We thank Iliana Samara for her help with data collection. This work was supported by a fellowship for postdoctoral researchers funded by the Alexander von Humboldt Foundation (to RLvdB), funding from the German Research Foundation (DFG; grant numbers DO 1240/3-1, DO 1240/4-1, and SFB 936/A7, to Tobias Donner), an FWO [PEGASUS]^2^ Marie Skłodowska-Curie fellowship (12T9717N, to KD), and a Consolidator Grant of the European Research Council (grant number GA 283314-NOREPI, to SN).

## Notes

**CONFLICT OF INTERESTS** The authors declare that no conflict of interests exists.

### Competing Interest Statement

The authors have declared no competing interest.

